# Short interval intracortical inhibition as measured by TMS-EEG

**DOI:** 10.1101/802504

**Authors:** Vishal Rawji, Isabella Kaczmarczyk, Lorenzo Rocchi, John C. Rothwell, Nikhil Sharma

## Abstract

The diagnosis of amyotrophic lateral sclerosis (ALS) relies on involvement of both upper (UMN) lower motor neurons (LMN). Yet, there remains no objective marker of UMN involvement, limiting early diagnosis of ALS. This study establishes whether TMS combined with EEG can be used to measure short-interval intracortical inhibition (SICI) via TMS evoked potentials (TEP) in healthy volunteers - an essential first step in developing an independent marker of UMN involvement in ALS.

We hypothesised that a SICI paradigm would result in characteristic changes in the TMS-evoked EEG potentials that directly mirror the changes in MEP.

TMS was delivered to the left motor cortex using single-pulse and three inhibitory stimulation paradigms. SICI was present in all three conditions. TEP peaks were reduced predominantly under the SICI 70 protocol but less so for SICI 80 and not at all for SICI 90. There was a significant negative correlation between MEPs and N45 TEP peak for SICI 70 (rho = −0.54, p = 0.04). In other words, as MEPs becomes inhibited the N45 increases. The same trend was maintained across SICI 80 and 90 (SICI 80, rho = −0.5, p = 0.06; SICI 90, rho = −0.48, p = 0.07). Additional experiments suggest these results cannot be explained by artefact.

We establish that motor cortical inhibition can be measured during a SICI 70 protocol expanding on previous work. We have carefully considered the role of artefact in TEPs and have taken a number of steps to show that artefact cannot explain these results and we suggesting the differences are cortical in origin. TMS-EEG has potential to aid early diagnosis and to further understand central and peripheral pathophysiology in MND.

## Introduction

Amyotrophic lateral sclerosis (ALS) is a universally fatal neurodegenerative disease without a cure or effective treatment. It is characterised by progressive degeneration of both upper (UMN) and lower motor neurons (LMN). While electromyography (EMG) directly assesses LMN function and conventional transcranial magnetic stimulation (TMS) depends on both UMN & LMNs, there remains no objective marker of UMN involvement. This directly limits early diagnosis of ALS and trial enrolment. This study establishes whether TMS combined with EEG can be used to measure short-interval intracortical inhibition (SICI) via TMS evoked potentials (TEP) in healthy volunteers - an essential first step in developing an independent marker of UMN involvement in ALS.

LMNs are affected in ALS therefore an independent marker of UMN is urgently required. TMS studies to date, that depend on both UMN & LMN, have demonstrated motor cortex dysfunction in ALS patients (1–3). In a patient series, Menon et al. show that SICI, can differentiate between patients with ALS and mimic disorders (1). However, patients with significant LMN involvement were excluded (specifically if a median nerve compound muscle action potential of <1 mV was not achieved or there was marked wasting of the thenar eminence - both of which are common in ALS patients). Despite these limitations, this important study supports the hypothesis of general hyperexcitability of the motor cortex in ALS.

Concurrent TMS-EEG of the motor cortex has the potential to measure UMN function without the confounds of LMN involvement. Using this technique, TMS stimulates the motor cortex and EEG acts as the readout, giving rise to TEPs. Critically, TEPs are devoid of peripheral confounds as they do not depend on a peripheral output. An MEP (the peripheral component) can be also measured simultaneously in healthy populations helping to establish the limitations of the method before application to patient populations. Concurrent TMS-EEG has potential as a diagnostic test in ALS by exclusively assessing motor cortex function.

In order to apply TMS-EEG to patient populations we first need to explore the protocol in healthy volunteers. Three previous studies have investigated SICI using TMS-EEG (4–6). In addition to the small numbers (n = 12, 8 and 16, respectively), only one set of preconditioning stimulus intensities has been used, resulting in a limited assessment of the method. It is widely accepted that changing the intensity of the preconditioning stimulus changes the degree of corticospinal inhibition (measured by MEPs) (7,8). However, it is not known how changing the preconditioning stimulus affects cortical inhibition, measured by TEPs. This is essential in understanding the relationship between TEP and MEP measured during a SICI protocol.

To understand the relationship between cortical and corticospinal inhibition we concurrently measured TEPs and MEPs, during SICI using a range of conditioning stimuli. We hypothesised that a SICI protocol will result in changes in the TMS-evoked EEG potentials that would reflect the changes in the MEP. More specifically, we hypothesised that cortical inhibition induced during SICI would be reflected within the TEP peaks (Experiment 1). It is accepted that there exists interindividual heterogeneity in the conditioning stimulus intensity required to produce maximal SICI. Consequently, group effects seen during SICI conditions may hide true cortical inhibition. That is, the effect of conditioning stimulus may be driving the differences in TEPs rather than reflecting true cortical inhibition. To account for this, we also calculated the average TEP waveform based on the degree of MEP inhibition (inhibition-focused analysis).

A recent report by Conde et al. suggests that despite state-of-the-art artefact removal techniques, the TEP includes significant, non-transcranial auditory and somatosensory artifacts (23). Given these concerns, we performed a number of steps to establish this neural origin of the SICI TEP signal in this study.

First, accepting that TMS artefacts on TEPs are a function of the TMS stimulator intensity, a SICI protocol with increasing conditioning stimuli intensities may result in artefact-based differences between the groups. To explore this, we ranked the data (from Experiment 1) based on the degree of MEP inhibition to test the hypothesis that differences between the SICI groups would effectively ‘disappear’ if they were solely a result of TMS stimulator intensity. Second, given TMS artefacts on TEPs are a function of TMS stimulator intensity, the artefact produced by individual pulses will largely be the same as during a paired pulse. In principle, subtracting the resulting TEP would produce a ‘flat’ waveform if it were a result of artifact. We hypothesised that subtraction of the single pulse TEPs from the paired pulse (SICI) TEP would result in no signal. In other words, we explore whether the resultant SICI TEP is artifact or neural in origin. Not only did we apply this to Experiment 1 but we also conducted a separate experiment (Experiment 2) to measure the TEP during SICI (at 70% of RMT followed after 2ms by 120% RMT) and from each single TMS pulse (at 70% of RMT & 120% RMT).

This study is important as it establishes whether concurrent TMS-EEG can measure SICI and the contribution - if any - of auditory and somatosensory artifacts before applying this to patient populations.

## Methods

### Participants

In Experiment 1, 15 right-handed subjects (10 female, mean age 24.07, SD 3.79) participated in this experiment. 8 subjects participated in Experiment 2 (right-handed subjects 4 female, mean age 23.38, SD 1.87). The study was approved by University College London Ethics Committee. Inclusion criteria included healthy, consenting adults over the age of 18 of either gender and right-handed. Exclusion criteria included implanted metal objects or devices (cochlear implant or deep brain stimulator) in the brain or skull, taking pro-epileptogenic medication or a history of spinal surgery. No subject had contraindications to TMS, which was assessed by a TMS screening questionnaire.

### Transcranial Magnetic Stimulation and Electromyography Recordings

Throughout the experiment, subjects were seated comfortably in a non-reclining chair, with their forearms were supported using a cushion. Electromyographic (EMG) activity was recorded from the right, first dorsal interosseous (FDI) muscle using 19 mm x 38 mm surface electrodes (Ambu WhiteSensor 40713) arranged in a belly tendon montage. The raw signals were amplified, and a bandpass filter was also applied (20 Hz to 2 kHz (Digitimer, Welwyn Garden City, United Kingdom)). Signals were digitised at 5 kHz (CED Power 1401; Cambridge Electronic Design, Cambridge, United Kingdom) and data were stored on a computer for offline analysis (Signal version 5.10, Cambridge Electronic Design, United Kingdom).

Single pulse, monophasic TMS was employed using a Magstim 200^2^ stimulator (The Magstim Co. Ltd) connected via a figure-of-eight coil with an internal wing diameter of 70 mm. A 0.5 cm foam layer was placed underneath the coil to minimise bone conduction of the TMS click and scalp sensation caused by coil vibration. The hotspot was identified as the area on the scalp where the largest and most stable motor evoked potentials (MEPs) could be obtained for the right first dorsal interosseous (FDI) muscle, using a given suprathreshold intensity. The coil was held approximately perpendicular to the presumed left central sulcus and tangentially to the skull, with the coil handle pointing backwards for postero-anterior (PA) stimulation. The resting motor threshold (RMT) was then found. This was defined as the lowest TMS stimulus intensity to evoke a response of 50 μV in 5 out of 10 trials in the relaxed FDI using the optimal PA orientation.

Single-pulse TMS was delivered at an intensity required to evoke a motor-evoked potential (MEP) of 1 mV. This served as a measure of unconditioned corticospinal excitability. To measure SICI in experiment 1, we delivered a preconditioning pulse 2 ms before single-pulse TMS, the intensity of which changed as a proportion of the RMT (70, 80, 90% of RMT) (9,10). Hence, we had four TMS conditions: spTMS (to measure unconditioned corticospinal excitability) and three SICI paradigms (SICI 70, SICI 80 and SICI 90) to measure corticospinal inhibition. 20 pulses at each condition were delivered per block, for four blocks. resulting in a total of 320 pulses per session. The peak-to-peak amplitude of MEPs served as a measure of corticospinal excitability. Consequently, the decrease in MEP amplitude under SICI conditions gives an indication of corticospinal inhibition.

For experiment 2 we then explored the individual components of the paired-pulse signal further. In experiment 2, we included four stimulating conditions: spTMS at 70% RMT, spTMS at 120% RMT, paired-pulse TMS (70% RMT, 70% RMT) and paired-pulse TMS (70% RMT, 120% RMT). Stimuli were delivered 2 ms apart from each other in the paired-pulse TMS conditions.

### Electroencephalographic recordings

EEG was simultaneously recorded with TMS using the 64-channel actiChamp System (Brain Products GmbH, Gilching, Germany) in accordance with the 10-20 international EEG electrode array. EEG signals were sampled at 5 kHz and impedances were kept below 5 kΩ. Recordings were referenced to the O_z_ electrode and FP_z_ was made the ground electrode. During online recordings, participants wore earphones covered with headphones, which continuously played white noise. The intensity of this noise was titrated up by the subject to mask the sound of the TMS click and hence reduce any TMS-evoked auditory potentials (11).

TEPs were processed in MATLAB (Version 2017a, MathWorks Inc., Natick, USA) using EEGLAB (12) in conjunction with a series of functions from the TMS-EEG signal analyser (TESA) toolbox (13). The TESA toolbox is a specific set of functions designed to analyse TEPs.

Preprocessing was in accordance with the established analysis protocol by Rogasch et al. (13) All trials, regardless of trial type, were preprocessed together and then separated into their corresponding types (experiment 1: spTMS, SICI 70, SICI 80 and SICI 90; experiment 2: 70% RMT, 120% RMT, 70% RMT x 70% RMT, and 70% RMT x 120% RMT). EEG signals were epoched (± 1.3 seconds) around the test pulse and demeaned from −1000 to - 10 ms before the test pulse. Individual trials were visually inspected and rejected if they were particularly noisy. The TMS artefact was then removed from −5 to 15 ms after the time of stimulation, interpolated and then the signal was downsampled to 1000 Hz. A first round of independent component analysis (ICA) using FastICA (14,15) was then performed to remove large, TMS-evoked muscle artefacts, after which band-pass (1-100 Hz) and band-stop (48-52 Hz) fourth order Butterworth filters were applied. Epoch length was reduced to ± 1 second to exclude possible edge artefacts due to filtering. A second round of ICA was then applied to remove residual artefacts based on time, frequency, scalp distribution and amplitude criteria described by Rogasch et al. (13) Finally, trials were re-referenced to the average reference and binned according to their corresponding trial type for post-processing.

Local motor cortical excitability was measured by averaging the TEP waveform in a cluster of four electrodes around the site of stimulation (C1, C3, CP1, CP3). This was performed for each condition and the amplitude of characteristic motor TEP peaks (N15, P30, N45, P60, N100 and P180) was calculated (5,16–21).

### Data analysis

Preprocessing of the EEG data was performed in Matlab using the EEGLAB toolbox (12). Statistical analysis was performed in R (Version 3.5.2).

#### Experiment 1

##### Measuring corticospinal inhibition during SICI

In keeping with the typical approach used in SICI studies, for each subject, we averaged individual MEPs for each condition, giving an overall indication of corticospinal excitability for that condition. As we were interested in confirming whether we had achieved corticospinal inhibition in our SICI conditions and hence performed Wilcoxon signed-rank tests between each SICI condition and spTMS.

##### Measuring cortical inhibition during SICI

After TMS to the motor cortex, characteristic peaks emerge – N15, P30, N45, P60, N100 and P180, which are believed to reflect physiological consequences of motor cortex stimulation. Based upon our hypothesis, we calculated the average TEP waveform for the four electrodes around the motor cortex (C1, C3, CP1, CP3), for each stimulating condition in each subject. We extracted the corresponding TEP peak amplitudes from the motor cortex TEP by finding the peak TEP amplitude in a prespecified window surrounding each time point. For example, the N15 peak was found by finding the negative peak of the TEP between 0 and 20 ms after TMS. For P30 this was between 15 and 35 ms, N45 between 30 and 55 ms, P60 between 50 and 70 ms, N100 between 90 and 150 ms, and P180 between 150 and 250 ms. The polarity of the peak found was in accordance with whether it was a positive or negative defined peak. If no peak was found, then we used the value at that time point (e.g. if no P30 peak was found, the TEP amplitude at 30 ms was used).

As with analysis of corticospinal inhibition using MEPs, we were interested in the changes in TEP peaks between our SICI conditions and spTMS. To assess changes in cortical inhibition induced by SICI compared to spTMS, we performed Wilcoxon signed-rank tests between each spTMS TEP peak (N15, P30, N45, P60, N100 and P180) and each corresponding SICI condition TEP peak. Non-parametric tests were used due to the distribution of the data.

We plotted the mean TEP waveform for all subjects (n=15), for spTMS and SICI TMS conditions. Making no assumptions on significance of TEP peaks, we performed a Wilcoxon signed-rank test for timepoints from TMS up to 200 ms after stimulation, whilst correcting for multiple comparisons using the false-discovery rate (FDR) correction method.

#### The relationship between the MEP and TEP

To further explore the relationship between MEPs and TEPs, we performed Spearman’s rank correlation between MEPs and TEP peaks for each SICI condition.

#### Exploring for artifact

Accepting that TMS artefacts on TEPs are a function of the TMS stimulator intensity, a SICI protocol with increasing conditioning stimuli may result in artefact-based differences between the groups. To explore this, we ranked the data (from Experiment 1) based on the degree of MEP inhibition. In other words, the intensity of the preconditioning stimulus was distributed across the groups. These MEPs were split into three equal groups according to the degree of inhibition (high, medium, low) and their corresponding TEP waveform analysed by performing Wilcoxon signed-rank tests for timepoints from TMS up to 200 ms after stimulation (with FDR correction). We predicted that if the differences between spTMS TEPs and SICI 70, 80 or 90 TEPs were a function of the TMS stimulator intensity, there would be no difference between the high, medium and low groups.

To further examine for artefact, we then subtracted the TEP waveform induced by SICI 70 (70% RMT & simulator output to produce a 1mV MEP) from that induced by spTMS (stimulator output to produce a 1mV MEP). With the assumption that the auditory and somatosensory artefacts induced by these two paradigms are broadly equivalent, the resultant waveform from mathematical subtraction should be zero if it is a result of artifact. We performed a one sample t-test on this resultant waveform to assess whether it significantly deviated from electroneutrality.

We further explored the possible effect of artefact on the TEP in an additional experiment **(Experiment 2)**. In principle, comparing the TEP from a single pulse TMS stimulus to a SICI paradigm involving the same TMS pulses should invoke that same artifact. To this end, we compared TEPs from two conditions (spTMS 70% RMT vs paired-pulse, where both stimuli were given at 70% RMT with the same interval of 2 ms). The resultant TEP waveforms were compared using Wilcoxon signed-rank tests with FDR correction.

Finally, it is feasible that a smaller TEP after SICI TMS, compared to spTMS, could reflect a refractory process of the neurons activated by the preconditioning stimulus. That is, the test stimulus is unable to activate the neuronal population that was activated by the preconditioning stimulus. If this was the case, then the difference in TEP amplitude between spTMS (120% RMT) and SICI TMS (70% RMT x 120% RMT) would be equivalent to the TEP evoked by the preconditioning stimulus alone (70% RMT). We therefore compared the spTMS 70% RMT TEP waveform and a TEP waveform made from the subtraction of SICI (70% RMT x 120% RMT) from spTMS 120%RMT alone.

## Results

### Physiological measurements

Mean resting motor threshold and 1 mV intensity was measured at 53.9% (SD 8.2) and 64.4% (SD 9.0) of the maximum stimulator output, respectively.

### The effect of SICI on MEPs

In keeping with the wider literature, we found that SICI decreased corticospinal excitability. Wilcoxon signed-rank tests showed that corticospinal excitability was significantly lower than spTMS for SICI 70 (p = 0.003), SICI 80 (p = 0.001) and SICI 90 (p = 0.005). These results confirmed that corticospinal excitability was suppressed during each of our SICI protocols and are shown in **Figure 1A**.

**Figure 1:**
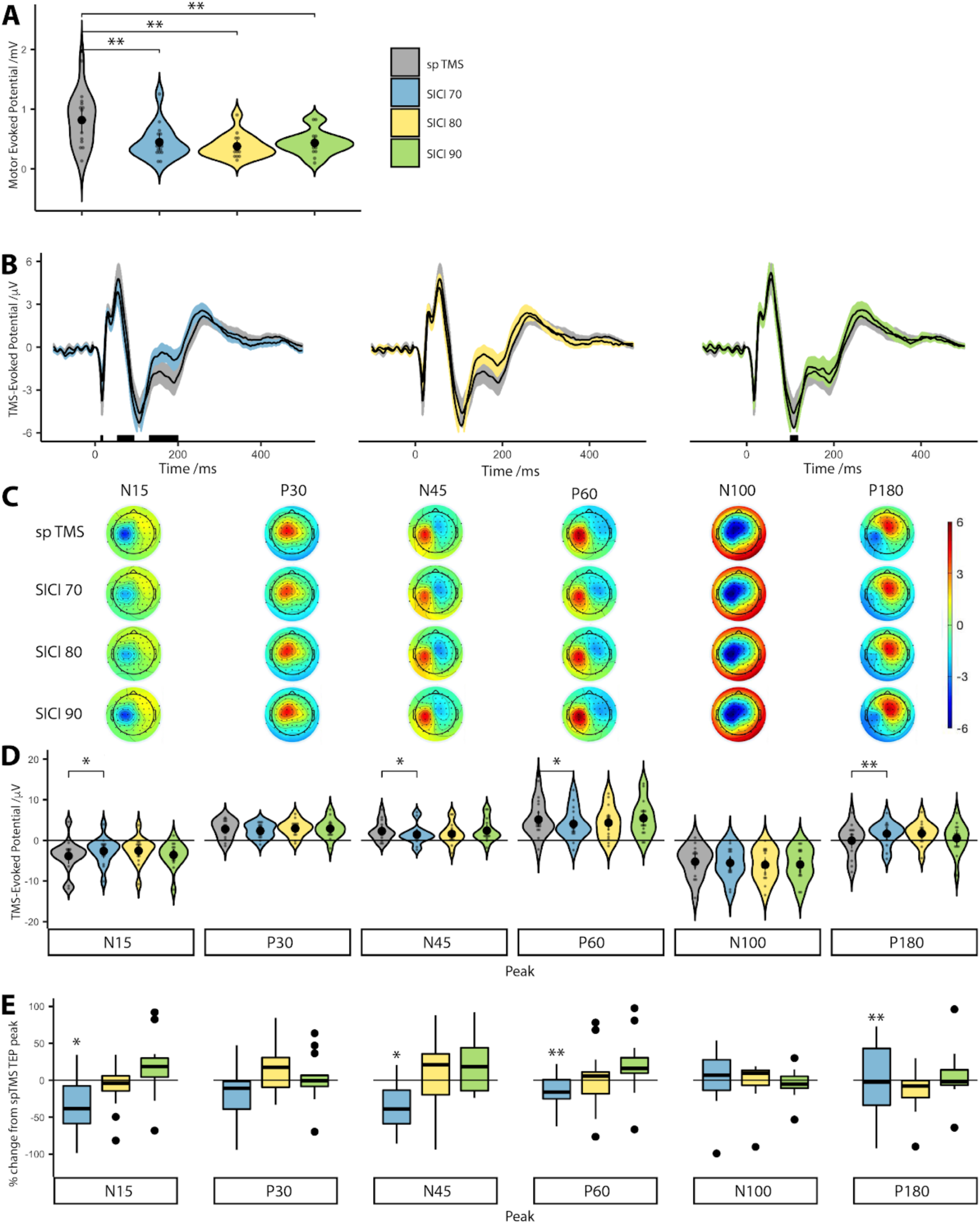
Corticospinal and cortical inhibition measured during SICI protocols. A: Corticospinal excitability measured by MEP amplitudes after spTMS and each of the SICI conditions. B: Average TEP waveform from four electrodes around the site of stimulation (C1, C3, CP1 and CP3). Shaded areas around the TEP waveform represent standard error of the mean. Black bars under each plot represent statistically significant (p < 0.05) time points, calculated from Wilcoxon signed-rank tests, after FDR correction. C: Scalp plots showing the TEP distribution across our six timepoints of interest, for each condition. D: TEP peaks at six time-points (15, 30, 45, 60, 100 and 180 ms) are extracted for each subject and plotted for each stimulation condition. E: Percentage change of individual TEP peaks during SICI conditions with respect to spTMS TEPs. Due to differences in polarity between conditions, values were squared first and then represented as a fraction of the spTMS TEP peak. Values below the dashed line depict suppression; those above depict facilitation. Asterisks represent statistically significant differences (* = p < 0.05, ** = p < 0.01).

### The effect of SICI on TEPs

We assessed cortical excitability measured using TEPs after spTMS and SICI. Making no assumptions on the significance of peaks, Wilcoxon signed-rank tests, with FDR correction, showed that only the SICI 70 TEP waveform differed significantly from the spTMS TEP waveform (**Figure 1B**).

Wilcoxon signed-rank tests during SICI 70 TMS indicated that the median SICI 70 TMS TEP test ranks were statistically different than the median spTMS TEP ranks for the N15 (Z = 23, p = 0.035), N45 (Z = 101, p = 0.018), P60 (Z = 105, p = 0.008) and P180 (Z = 11, p = 0.003) peaks. The P30 (Z = 69, p = 0.639) and N100 (Z = 73, p = 0.489) peak ranks did not differ statistically from spTMS TEP peaks.

For SICI 80 there was a trend towards difference for N15 (Z = 28, p = 0.073) and N100 (Z = 90, p = 0.095 and P180: Z = 26, p = 0.055). There were no statistically significant ranks differences between the median ranks for the remaining peaks (P30: Z = 55, p = 0.804; N45: Z = 88, p = 0.121; P60: Z = 81, p = 0.252;).

For SICI 90, there was no statistically significant ranks differences between the median ranks (N15: Z = 58, p = 0.934; P30: Z = 51, p = 0.639; N45: Z = 63, p = 0.890; P60: Z = 57, p = 0.890; N100: Z = 84, p = 0.188 and P180: Z = 49, p = 0.561). As we included the amplitude even when a peak was not present, we re-analysed the data including only peaks and the results did not significantly alter.

We normalised SICI TEP peak amplitudes to spTMS conditions, to directly compare percentage changes of individual peak amplitudes at different SICI conditions (Figure 1E). In SICI 70 conditions, Wilcoxon signed-rank tests indicated a significant peak reduction in N15 (Z = 96, p = 0.041), N45 (Z = 19, p = 0.018), P60 (Z = 15, p = 0.008) and P180 (Z = 109, p = 0.003). There was a trend towards a reduction in peaks N15 (Z = 90, p = 0.073) and P180 (Z = 94, p = 0.055). There were no significant changes in remaining TEP peaks at SICI 80 (P30: Z = 65, p = 0.804; N45: Z = 32, p = 0.121; P60: Z = 39, p = 0.252; N100: Z = 30, p = 0.095), SICI 90 conditions did not show significant percentage changes in any of the TEP peaks (N15: Z = 58, p = 0.934; P30: Z =6 9, p = 0.639; N45: Z = 57, p = 0.891; P60: Z = 63, p = 0.981; N100: Z = 36, p = 0.188, P180: Z = 72, p = 0.525).

We conclude that despite MEP suppression for all three SICI conditions, the TEP did not follow the same pattern. TEP peaks were reduced predominantly under the SICI 70 protocol but less so for SICI 80 and not at all for SICI 90.

### Relationship between MEPs & TEPs

To explore the relationship between MEPs and TEPs, we performed Spearman’s rank correlation coefficients between MEPs and TEP peaks for each SICI condition. The results are shown in **Figure 2**.

**Figure 2:**
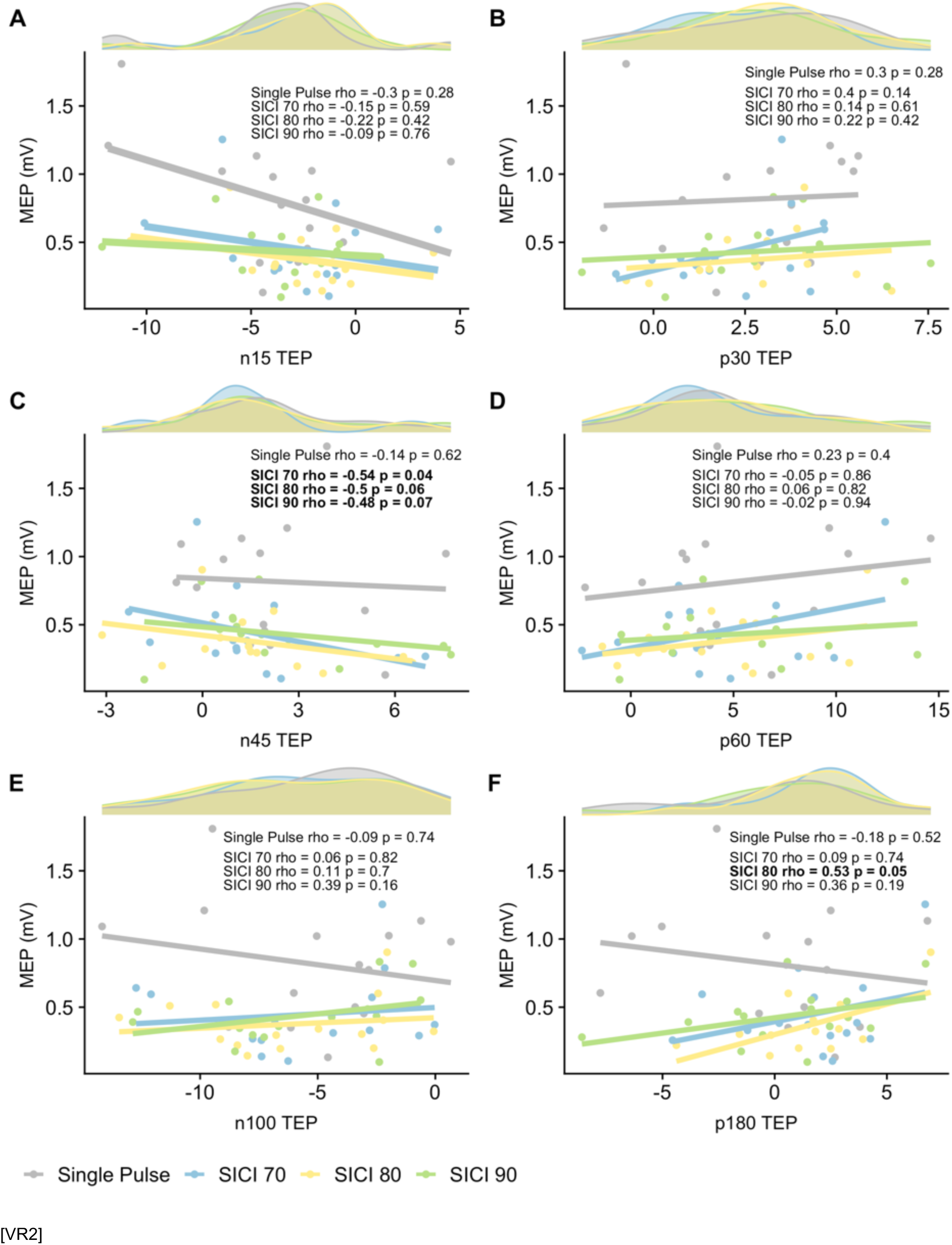
Spearman rank correlations between MEP and TEP peaks for single pulse, SICI 70, 80, 90. Each plot shows Spearman rank correlations between the MEP amplitude and TEP peak amplitude (N15, P30, N45, P60, N100 and P180), for spTMS and each SICI condition. Above each plot is the distribution of each peak per condition.

There was a significant negative correlation between MEPs and N45 TEP peak for SICI 70 (rho = −0.54, p = 0.04). In other words, as MEPs becomes inhibited the N45 increases. The same trend is maintained across SICI 80 and 90 (SICI 80, rho = −0.5, p = 0.06; SICI 90, rho = −0.48, p = 0.07).

There was a positive correlation between MEPs and P180 for SICI 80 only (rho = 0.53, p = 0.05).

#### Exploring artefact

As TMS artefacts on TEPs are a function of the TMS stimulator intensity, a SICI protocol with increasing conditioning stimuli may result in artefact-based differences between the groups. The individual SICI derived MEPs were ranked according to their peak-to-peak amplitude, regardless of which SICI condition they arose from. These MEPs were split into three groups according to the degree of inhibition (high, medium, low) and their corresponding TEP waveform analysed. This demonstrated significant inhibition of the TEP in the high inhibition group (when compared to spTMS TEPs) suggesting that the effect of a SICI protocol on TEPs are not a result of the stimulator output. This also shows that when accounting for interindividual differences in conditioning stimulus intensity required to produce maximal SICI, TEPs still show differences when MEPs are maximally suppressed. This confirms that cortical inhibition can indeed be measured using TEPs.

**Figure 3** shows the subtraction of the spTMS (1mV TEP) and SICI 70 TEP waveforms. A one sample t-test with FDR correction performed on the resultant waveform shows that the TEP significantly differs from zero, indicating that when controlled for non-transcranial, multisensory artefacts, SICI 70 TMS induces a neural signal as measured by the TEP. The findings were similar in Experiment 2 when subtracting spTMS 70% RMT and paired-pulse TMS 70% 70% RMT, as well as when subtracting SICI TEPs (70% RMT x 120% RMT) from spTMS TEP (120% RMT).

**Figure 3:**
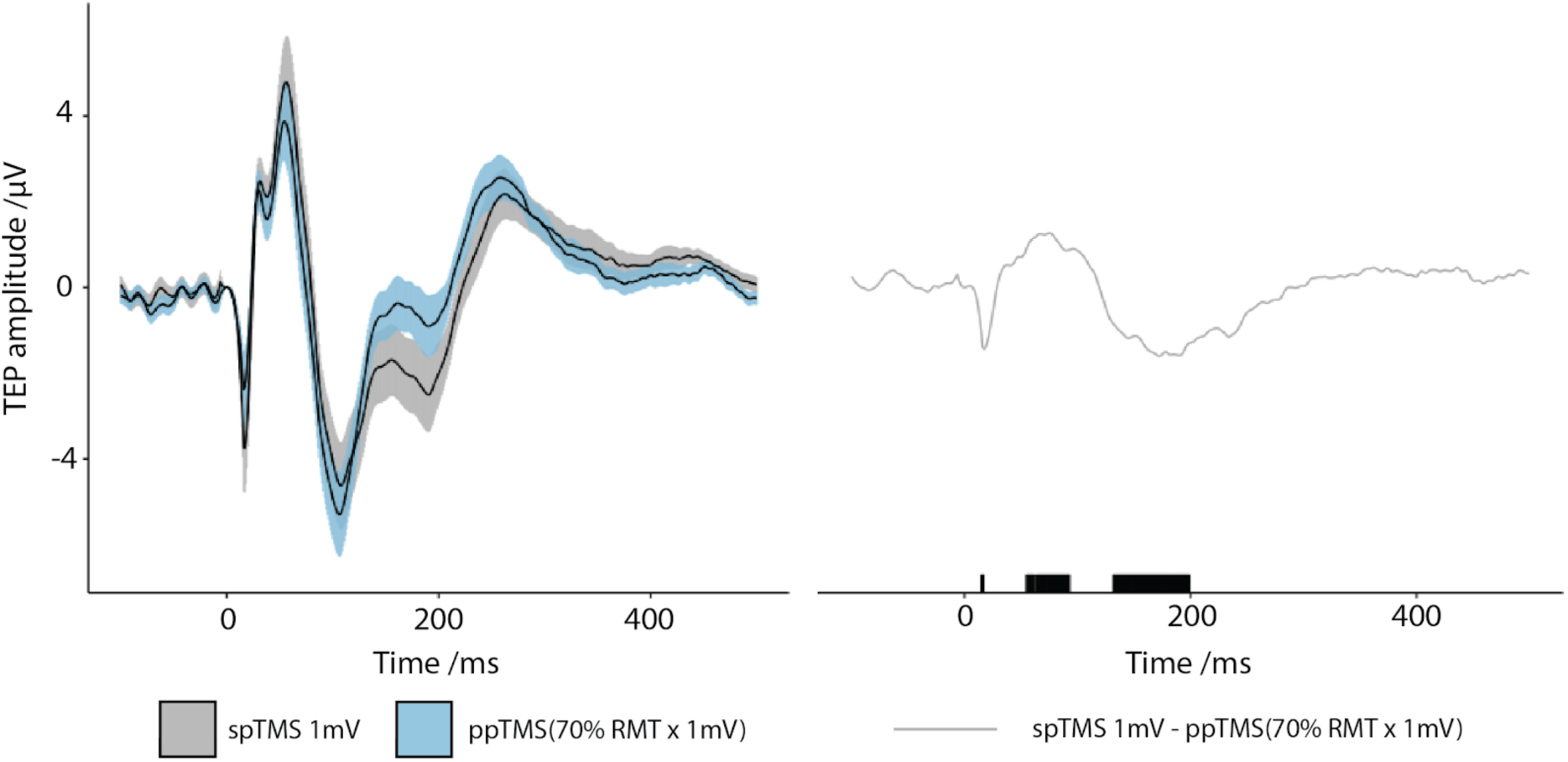
Subtracted TEP waveform for SICI 70. Top plot shows the TEP waveform for SICI 70 and spTMS. Bottom plot shows the resulting TEP waveform after mathematical subtraction (spTMS – SICI 70), with black bars indicating regions of statistical significance (p < 0.05) after one sample t-test FDR correction.

We considered that the decrease in SICI TEP amplitude found in Experiment 1 may be due to a refractory period set up by the preconditioning stimulus. If true, then subtraction of SICI TEPs from spTMS TEPs should be equal to the preconditioning stimulus TEP. Subtraction of SICI (70% RMT x 120% RMT) from spTMS (120% RMT) did not result in the same waveform as 70% RMT TEP, thereby suggesting that the decrement in SICI TEP amplitude was due to an inhibitory process rather than a refractory period set up by the preconditioning stimulus.

## Discussion

In this proof of principle study in healthy volunteers, we established that motor cortical inhibition can be measured during a SICI protocol expanding on previous work (4–6). Our results show that cortical inhibition was achieved at SICI 70. We have carefully considered the role of artefact in TEPs and have taken a number of steps to show that artefact cannot explain these results suggesting the differences are cortical in origin. This includes removing dose dependent artifact during the SICI paradigm (by re-analysing the data after ranking by corticospinal inhibition) and subtraction of the SICI TEP from the relevant single pulse TEP in two different experiments. This study highlights the similarities but also important differences between the TEP peaks and MEP inhibition during a SICI protocol that need to be considered carefully when applying this to conditions such as MND.

We report that during SICI 70 TMS, there is significant inhibition of the N15, N45, P60 and P180 TEP peaks. Our findings are supported by previous literature, particularly modulation of the N45 (4–6), which was significantly correlated with the degree of corticospinal inhibition. The earliest observable peak in our study, the N15, has previously been suggested to reflect activation of the posterior parietal cortex, following M1 TMS (5,24). In contrast, the N45 peak thought to be generated by the motor cortex as suggested by dipole modelling (19). We also find that the degree of N45 peak modulation scales with corticospinal excitability during SICI suggesting it could be a reflection of the motor cortex and/or spinal excitability. The P60 peak, which was also modulated, is thought to reflect general cortical excitability (4,17,25).

The interpretation of specific TEP peaks needs to be carefully considered. To date, pharmacologic manipulation (26–31) and correlation with measures of corticospinal excitability have been used to hypothesise the role of certain peaks. For example, the N100 peak being implicated in GABA_B_ergic transmission (32) and the P30 peak being related to cortical excitability (4,17,25). Furthermore, the N100-P180 complex has been shown to be significantly modulated during SICI compared to spTMS (6), although we found a statistically significant difference of only the P180 peak during SICI. Hence, the peaks modulated during SICI should conservatively be regarded as epiphenomena of cortical inhibition. TEPs have been considered to represent sensory feedback from muscle activation, which may contaminate TEP measurements. Whilst this is a caveat to the interpretation of TEPs, the latency of any sensory feedback would realistically affect those later peaks such as the P60, although the event-related desynchronisation, believed to reflect afferent proprioceptive information, has been observed after 300 ms post-stimulation (33). Nevertheless, the findings of the N15, P30 and N45 peaks should remain unaffected by the confound of sensory feedback.

TEP peaks convincingly demonstrate cortical inhibition during a standard SICI protocol. This was statistically significant for the SICI 70 condition. This is in contrast to the SICI 80 and 90 conditions where the cortical inhibition did not reach significance, contrasting with the simultaneously measured MEPs. There are a number of possible explanations for this. The approach to analysis and underlying assumptions of MEPs and TEPs are very different, which may explain the discrepancy between corticospinal and cortical inhibition. MEP analysis requires few steps, with peak-to-peak amplitude measurements of an amplified signal using surface EMG electrodes. Conversely, TEP analysis has several steps such as filtering, noise reduction and independent component analysis. While, we followed the steps for a previously established TEP analysis pipeline (13,34), each step has a number of assumptions that manipulates the data. While the underlying process may be the same, the difference in the approaches may alter the output and in effect ‘hide’ the effects of cortical inhibition from TEP analysis. We do not believe this is simply a result of being underpowered as we were able to measure cortical inhibition for the SICI 70 condition.

Alternately, a key difference between SICI 70, 80 and 90 is the greater stimulator output that may have a differential effect on the EEG electrodes compared to the surface EMG. Surface EMG is remote from the TMS coil limiting local effects unlike the EEG electrodes. Indeed, further interrogation of this discrepancy between corticospinal and cortical inhibition revealed that this may be due to the effect of the preconditioning stimulus. As we predicted, TEP peaks were more strongly correlated with MEP size for SICI conditions with smaller preconditioning stimuli. This supports the hypothesis that increasing the preconditioning stimulus results in additional noise in the EEG signal that is not reflected in the MEP.

Another possible explanation is that, unlike EEG, surface EMG data represents a highly filtered signal. The EEG electrode configuration uses 64 electrodes to record cortical activity. TEPs represents a composite signal of cortico-cortical activity whereas MEPs measure corticospinal activity for a signal muscle (33). TMS recruits a heterogenous mixture of neural elements, including excitatory and inhibitory interneurons, and even interneurons not relevant to the FDI (35); EEG does not differentiate between these different sources, whereas the FDI MEP represents a somatotopically sensitive output. In other words, the FDI MEP represents a highly filtered signal. In fact, the MEP is a smoothed average of descending cortical output onto spinal motor neurons (36). The motor cortex TEP, on the other hand, is an unfiltered signal (is less topographically selective for the FDI) comprised of activity relevant to the FDI cortical motor neurons and many other neurons that are not our primary focus. Consequently, increasing preconditioning stimulus intensity increasingly results in activation of neurons not relevant to the FDI axis and this extra neural noise (i.e. not FDI relevant) contributes to the TEP but not the MEP. Hence, whilst the FDI MEP may be inhibited, the TEP does not discriminate FDI related cortical activity from non-FDI related cortical activity. With increasing preconditioning stimulus intensities, the amount of this extra neural noise increases and the discrimination between spTMS and SICI TEPs, decreases. We propose that this is why SICI 70 TEPs are statistically different to spTMS TEPs, whereas SICI 80 and 90 are not. Going forward, we recommend that the intensity of preconditioning stimuli should be minimised when attempting to measure cortical inhibition using TMS-EEG.

To date, it has not been possible to investigate the interaction of paired-pulse TMS using a subthreshold conditioning *and* a subthreshold test stimulus as MEP’s require a suprathreshold stimulus to be measured. To our knowledge, paired-pulse TMS-EEG studies have also not investigated the interaction between subthreshold stimuli. We find that during 70% RMT x 70% RMT stimulation, the paired-pulse paradigm results in a TEP with greater amplitude than spTMS at 70% RMT. This needs to be interpreted with care as this experiment was designed to examine for artefacts. Models of SICI have suggested that the preconditioning stimulus results in a refractory period of low-threshold GABA-ergic neurons (37), which might explain the decrement in the TEP during SICI conditions. Future work will specifically address SICI with a subthreshold test stimulus pulse.

Despite state-of-the-art artefact removal techniques, a significant proportion of the TEP waveform has been attributed to multisensory, non-transcranial stimulation, predominantly from somatosensory, auditory and reafferent sources (23). Our experimental design inherently accounts for these artefacts; the artefacts induced by paired-stimuli separated by 2 ms reflect the artefacts evoked by suprathreshold stimuli. Hence, the subtraction of TEP waveforms accounts for the non-transcranial artefacts cited in (23) and reveals the underlying signal is neural in origin.

If the smaller TEP was due to a refractory population not being stimulated by the test stimulus, then the magnitude of this difference should be equal to the magnitude of the refractory population alone (conditioning stimulus TEP). Our finding that the subtraction of the test stimulus TEP from the SICI TEP is not equivalent to the preconditioning stimulus TEP alone shows that decrement in the TEP is not simply due to a refractory period of the neurons stimulated by the preconditioning stimulus. Instead, it provides further evidence (8,39) that SICI is produced via an inhibitory intracortical interaction.

This works lays the foundation to apply this method to patient populations although there remain a number of unanswered questions. For instance, work needs to explore the effect of ageing on SICI measured with TEPs (ALS typically affects older individuals). However, it should be noted that this may be difficult to resolve given the literature on SICI is more widely studied and yet remains unclear (40). Regardless, those studies assess corticospinal inhibition, as measured by the MEP; it may be the case that cortical inhibition measured by TEPs varies differently with aging. These findings encourage an assessment of motor cortex SICI during ageing.

This work is relevant for TMS SICI in ALS patients where it has been proposed as a tool to aid the diagnostic pathway. As highlighted above, SICI MEPs can be dependent upon factors other than cortical involvement. This strengthens the rationale for using TMS-EEG in ALS. More specifically the proposal is that measurement of corticospinal inhibition might be useful in diagnosing ALS (1). However, this is limited for the reasons stated in the Introduction. Furthermore, in the study by Menon et al. patients had to exhibit six months disease progression. In the context of ALS, six months to come to a diagnosis may now be considered to be too long to be of clinical use. Preferably, a diagnostic test in this context would be applied as early as possible, potentially alongside initial neurophysiological tests; it is unknown whether the changes in corticospinal inhibition occur in patients living with ALS early in the disease course. The major limitation that the MEP is a composite measure of UMN and LMN function, further undermines the use of a pure TMS protocol as a diagnostic tool.

## Conclusions

This work supports that a TMS-EEG approach to SICI may overcome many of the limitations faced by an exclusive MEP based approach and SICI 70 appears optimal. First and foremost, it avoids the limitations posed by LMN involvement in ALS. Consequently, patients with marked wasting can still be included in testing. Of course, in some patients it may be that MEPs can still be recorded during TMS-EEG and hence the benefits afforded by a TMS approach still apply during TMS-EEG - critically this would allow it to potentially become a powerful tool for longitudinal measurements to track progression.

